# Rapid mortality in captive bush dogs (*Speothos venaticus*) caused by influenza A of avian origin (H5N1) at a wildlife collection in the United Kingdom

**DOI:** 10.1101/2024.04.18.590032

**Authors:** Marco Falchieri, Scott M. Reid, Akbar Dastderji, Jonathan Cracknell, Caroline J. Warren, Benjamin C. Mollett, Jacob Peers-Dent, Audra-Lynne D Schlachter, Natalie Mcginn, Richard Hepple, Saumya Thomas, Susan Ridout, Jen Quayle, Romain Pizzi, Alejandro Núñez, Alexander M. P. Byrne, Joe James, Ashley C. Banyard

**Affiliations:** Influenza and Avian Virology Team, Department of Virology, Animal and Plant Health Agency (APHA-Weybridge), Woodham Lane, Addlestone, Surrey KT15 3NB, UK; Mammalian Virology Investigation Unit, Department of Virology, Animal and Plant Health Agency (APHA-Weybridge), Woodham Lane, Addlestone, Surrey KT15 3NB, UK; Knowsley Safari, Prescot, Merseyside L34 4AN, UK; Department of Pathology and Animal Sciences, Animal and Plant Health Agency (APHA-Weybridge), Woodham Lane, Addlestone, Surrey KT15 3NB, UK; APHA Field Epidemiology team, APHA Bridgwater, Rivers House, East Quay, Bridgwater, TA6 4YS; APHA Field Epidemiology team, APHA Hornbeam House, Electra Way, Crewe, Cheshire, CW1 6GJ; WOAH/FAO International Reference Laboratory for Avian Influenza, Animal and Plant Health Agency (APHA-Weybridge), Woodham Lane, Addlestone, Surrey KT15 3NB, UK; Worldwide Influenza Centre, The Francis Crick Institute, Midland Road, London. NW1 1AT

**Keywords:** Avian influenza, systemic infection, bush dogs, conservation species, terrestrial carnivores, H5N1, high pathogenicity, zoonotic assessment

## Abstract

Europe has suffered unprecedented epizootics of high pathogenicity avian influenza (HPAI) clade 2.3.4.4b H5N1 since Autumn 2021. As well as impacting upon commercial and wild avian species, the virus has also infected mammalian species more than ever observed previously. Mammalian species involved in spill over events have primarily been scavenging terrestrial carnivores and farmed mammalian species although marine mammals have also been affected. Alongside reports of detections in mammalian species found dead through different surveillance schemes, several mass mortality events have been reported in farmed and wild animals. During November 2022, an unusual mortality event was reported in captive bush dogs (*Speothos venaticus*) with clade 2.3.4.4b H5N1 HPAIV of avian origin being the causative agent. The event involved an enclosure of fifteen bush dogs, ten of which succumbed during a nine-day period with some dogs exhibiting neurological disease. Ingestion of infected meat is proposed as the most likely infection route.

## Introduction

High pathogenicity avian influenza virus (HPAIV) has caused significant mortalities across avian species since the emergence of H5Nx clade 2.3.4.4 viruses in Europe and the United Kingdom (UK) in 2014 (1, 2). In recent years, these viruses have undergone rounds of genetic diversification and caused repeated detections of H5Nx HPAIVs in poultry and wild birds, with each epizootic following a seasonal pattern. In Autumn 2021 the situation escalated significantly with clade 2.3.4.4b HPAIV H5N1 emerging to cause global outbreaks at a level never seen before (3, 4). Between 2014 and 2020, spillover of avian influenza infection into mammals was uncommon although occasional reports in both wild and captive mammals have been described (5, 6). With the escalation of H5N1 outbreaks during 2020/21 (3, 7), through the summer of 2021 (8) and into 2022/23 (4, 7, 9), infection pressure in the environment, as a result of high mortality levels in wild birds has led to an increase in the occurrence of spill over events globally (5, 10–18). To date, almost all detections in mammals have involved wild scavenging species that have most likely become infected following the ingestion of infected wild bird carcasses in the environment. These events have affected a broad range of species of both terrestrial and marine habitats (9, 19). Importantly, with the vast majority of cases the infection was reported in a single mammal following an undefined exposure to infectious material. However, on occasion, mammalian infection with avian origin H5N1 HPAIV has been implicated as the cause of mass mortalities. This has occurred most notably in natural events whereby wild animals have been affected (e.g., seals in the United States and sea lions in Peru (20, 21)) as well as in farmed species (farmed mink in Spain (22), farmed foxes in Finland (23)) and domesticated species (cats in Poland (24, 25) and South Korea (26)). These events have triggered interest in the potential for mammal-to-mammal transmission and in any resultant genetic adaptation. Assessment of transmission is undertaken both through epidemiological investigation and by thorough phylogenetic analysis with subsequent interrogation for potential adaptive mutations that may confer a replicative advantage to the novel mammal host (27). Whilst these outbreaks have shown concurrent infection of multiple individuals with the same virus, data supporting active transmission between mammals is lacking and a definitive conclusion on the ability of these viruses to spread directly from mammal to mammal has not been reached. On top of mortality events within the veterinary sector, this clade 2.3.4.4b H5N1 HPAIV has also been associated with human infection, with outcomes varying from asymptomatic to severe with hospitalisation of infected individuals (19, 28–34). These features have raised the zoonotic profile of the currently circulating H5N1 clade 2.3.4.4b HPAIV, although again human-to-human transmission has not been demonstrated.

Here we report on the infection and severe mortality within a pack of bush dogs (*Speothos venaticus*) in captivity with avian origin H5N1 clade 2.3.4.4b HPAIV. Bush dogs are a near threatened species of wild canids that are of conservation concern. Wild populations of these dogs range from northern regions of Panama (Central America) to northeastern Argentina and Paraguay; with populations also being present in Colombia, Venezuela, the Guianas, Brazil, and eastern Bolivia and Peru. This species is characterized by its small size, elongated body, small eyes, short snout, short tail, short legs, and small and rounded ears, in addition to gregarious and diurnal behaviour (35). In this disease event which occurred in November 2022, two thirds of the pack of bush dogs, held captive in a wildlife collection the UK, became clinically unwell with a disease that had a short duration and led to death and/or the need for euthanasia on welfare grounds with a range of clinical signs including neurological disease. Avian influenza was not suspected at first and several tests and analysis were performed at private laboratories to ascertain cause of death and to exclude the involvement of more common canine pathogens. Overall inconclusive results led bush dog samples to be submitted retrospectively to the Animal and Plant Health Agency (APHA), Virology Department for shotgun metagenomic assessment, which detected presence of influenza type A virus sequences in internal organs. We describe the disease event, timeline, virological and pathological impact of disease and sequence analysis of the causative agent.

## Methods

### Clinical investigation, post-mortem examination, tissue sampling and histopathological analysis

Clinical records including displayed signs and symptoms, blood chemistry data and mortality were recorded by the safari park personnel as part of the day-to-day activities. All bush dogs underwent full post-mortem examination (PME) at the index site, immediately after they were either euthanised or found dead. Of the ten animals, six were euthanised using intracardiac pentobarbitone (0.03 mg/kg, 1 mg/ml) under anaesthesia induced with medetomidine (0.03 mg/kg 1 mg/ml) and ketamine (4.03 - 4.85 mg/kg, 100 mg/ml), the remainder were found dead within the enclosure. Post-mortem examination was undertaken on site at the zoological collection with multiple tissues sampled for microbial culture, molecular analysis and formalin fixation. Tissue samples were submitted by the index site for bacteriological analysis to a private laboratory, frozen tissues were stored at -18 °C in a standard chest freezer for subsequent testing as required, while other tissues were stored in 10 % formalin and submitted for histology to a private laboratory pathologist. Later, remaining formalin fixed tissue, and formalin fixed paraffin embedded tissues from the private laboratory were further submitted to the Avian Influenza National Reference Laboratory at APHA for additional investigation (Table 1). Four-micron thick serial sections were stained with haematoxylin and eosin (H&E), and for immunohistochemistry (IHC), using a mouse monoclonal anti-influenza A nucleoprotein antibody (Staten’s Serum Institute, Copenhagen, Denmark) for histopathological examination and demonstration of influenza A virus nucleoprotein (NP). The overall distribution of virus-specific staining in each tissue was assessed using a semi-quantitative scoring system (where 0 = no staining, 1 = minimal, 2 = mild, 3 = moderate and 4 = widespread staining) modified from (36). Specificity of immunolabelling was assessed in positive control sections by replacing the primary antibody with a matching mouse IgG isotype; no non-specific cross-linking was observed. Frozen tissues for virological analysis (Table 2) were prepared as a 10% (w/v) suspension in Leibovitz’s L-15 medium and incubated at room temperature for 60 minutes before using standard RNA extraction protocols (37).

**Table 1:**
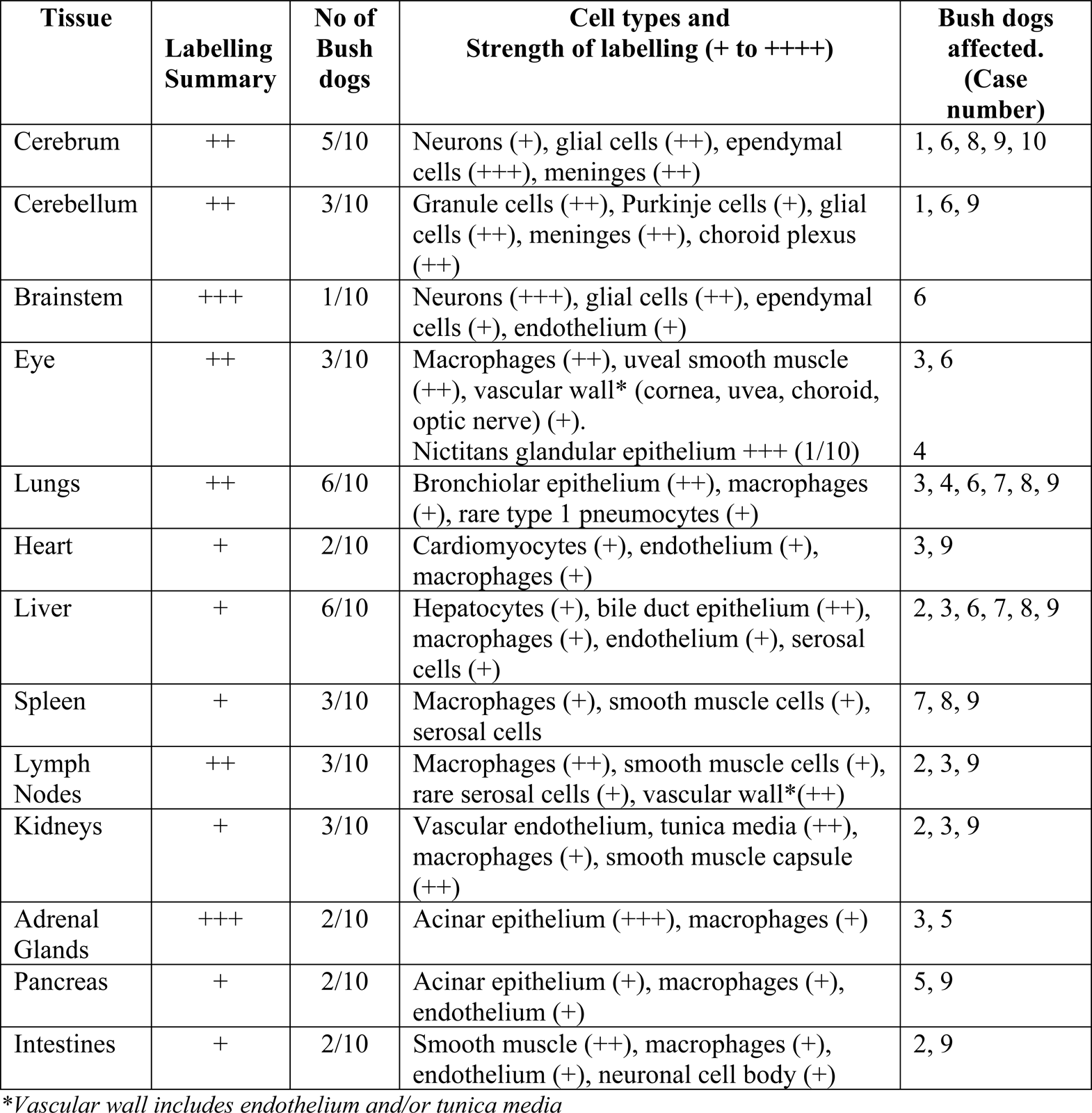
Immunohistochemistry findings in ten bush dogs infected with HPAI H5N1

| **Tissue** | **Labelling**<br>**Summary** | **No of**<br>**Bush**<br>**dogs** | **Cell types and**<br>**Strength of labelling (+ to ++++)** | **Bush dogs**<br>**affected.**<br>**(Case**<br>**number)** |
| --- | --- | --- | --- | --- |
| Cerebrum | ++ | 5/10 | Neurons (+), glial cells (++) , ependymal cells (+++), meninges (++) | 1, 6, 8, 9, 10 |
| Cerebellum | ++ | 3/10 | Granule cells (++) , Purkinje cells (+), glial cells (++) , meninges (++) , choroid plexus (++) | 1, 6, 9 |
| Brainstem | +++ | 1/10 | Neurons (+++), glial cells (++) , ependymal cells (+), endothelium (+) | 6 |
| Eye | ++ | 3/10 | Macrophages (++) , uveal smooth muscle (++) , vascular wall* (cornea, uvea, choroid, optic nerve) (+).<br>Nictitans glandular epithelium +++ (1/10) | 3, 6<br>4 |
| Lungs | ++ | 6/10 | Bronchiolar epithelium (++) , macrophages (+), rare type 1 pneumocytes (+) | 3, 4, 6, 7, 8, 9 |
| Heart | + | 2/10 | Cardiomyocytes (+), endothelium (+), macrophages (+) | 3, 9 |
| Liver | + | 6/10 | Hepatocytes (+), bile duct epithelium (++) , macrophages (+), endothelium (+), serosal cells (+) | 2, 3, 6, 7, 8, 9 |
| Spleen | + | 3/10 | Macrophages (+), smooth muscle cells (+), serosal cells | 7, 8, 9 |
| Lymph Nodes | ++ | 3/10 | Macrophages (++) , smooth muscle cells (+), rare serosal cells (+), vascular wall*(++) | 2, 3, 9 |
| Kidneys | + | 3/10 | Vascular endothelium, tunica media (++) , macrophages (+), smooth muscle capsule (++) | 2, 3, 9 |
| Adrenal Glands | +++ | 2/10 | Acinar epithelium (+++), macrophages (+) | 3, 5 |
| Pancreas | + | 2/10 | Acinar epithelium (+), macrophages (+), endothelium (+) | 5, 9 |
| Intestines | + | 2/10 | Smooth muscle (++) , macrophages (+), endothelium (+), neuronal cell body (+) | 2, 9 |

\*Vascular wall includes endothelium and/or tunica media

**Table 2:** Molecular assessment of post-mortem samples received for retrospective testing

| Case No. | Sample type | Sampling date | Real-time RT-PCR |  |  | Whole Genome Sequence (GISAID reference) |  |
| --- | --- | --- | --- | --- | --- | --- | --- |
|  |  |  | M-gene | HP H5 | N1 | Animal ID/Source | Accession Number |
| 1 | Liver | 17th Nov 2022 | + | + | + | / | / |
| 1 | Kidney | 17th Nov 2022 | + | + | + | / | / |
| 1 | Lung | 17th Nov 2022 | + | + | + | A/Bush dog/England/037675/2022 | EPI_ISL_17360134 |
| 1 | Spleen | 17th Nov 2022 | + | + | + | / | / |
| 2 | Ascitic fluid | 18th Nov 2022 | + | + | + | A/Bush dog/England/037680/2022 | EPI_ISL_17257532 |
| 2 | Viscera | 18th Nov 2022 | + | + | + | / | / |
| 3 | Ascitic fluid | 18th Nov 2022 | + | + | + | A/Bush dog/England/037682/2022 | EPI_ISL_17257533 |
| 3 | Intestine | 18th Nov 2022 | + | + | + | / | / |
| 4 | Ascitic fluid | 19th Nov 2022 | + | + | + | A/Bush dog/England/037684/2022 | EPI_ISL_17257534 |
| 4 | Intestine | 19th Nov 2022 | + | + | + | / | / |
| 4 | Liver | 19th Nov 2022 | + | + | + | / | / |
| 5 | Ascitic fluid | 19th Nov 2022 | + | + | + | / | / |
| 5 | Intestine | 19th Nov 2022 | + | + | + | / | / |
| 5 | Kidney | 19th Nov 2022 | + | + | + | / | / |
| 5 | Liver | 19th Nov 2022 | + | + | + | / | / |
| 5 | Urine | 19th Nov 2022 | + | + | + | A/Bush dog/England/068984/2022 | EPI_ISL_17710037 |
| 6 | Brain | 20th Nov 2022 | + | + | + | A/Bush dog/England/058198/2022 | EPI_ISL_17521050 |
| 6 | Intestine | 20th Nov 2022 | + | + | + | / | / |
| 6 | Kidney | 20th Nov 2022 | + | + | + | / | / |
| 6 | Liver | 20th Nov 2022 | + | + | + | / | / |
| 6 | Lung | 20th Nov 2022 | + | + | + | / | / |
| 6 | Oral swab | 20th Nov 2022 | NEG | NEG | NEG | / | / |
| 6 | Rectal swab | 20th Nov 2022 | NEG | NEG | NEG | / | / |
| 7 | Intestine | 21st Nov 2022 | + | + | + | A/Bush<br>dog/England/037689/2022 | EPI_ISL_17257535 |
| 7 | Kidney | 21st Nov 2022 | + | + | + | / | / |
| 7 | Liver | 21st Nov 2022 | + | + | + | / | / |
| 8 | Intestine | 21st Nov 2022 | + | + | + | A/Bush<br>dog/England/037690/2022 | EPI_ISL_17257536 |
| 8 | Kidney | 21st Nov 2022 | + | + | + | / | / |
| 8 | Liver | 21st Nov 2022 | + | + | + | / | / |
| 8 | Spleen | 21st Nov 2022 | + | + | + | / | / |
| 9 | Ascitic fluid | 22nd Nov 2022 | + | + | + | / | / |
| 9 | Brain | 22nd Nov 2022 | + | + | + | / | / |
| 9 | Intestine | 22nd Nov 2022 | + | + | + | / | / |
| 9 | Kidney | 22nd Nov 2022 | + | + | + | / | / |
| 9 | Liver | 22nd Nov 2022 | + | + | + | / | / |
| 9 | Lung | 22nd Nov 2022 | + | + | + | / | / |
| 9 | Pleura | 22nd Nov 2022 | + | + | + | A/Bush<br>dog/England/037691/2022 | EPI_ISL_17257537 |
| 9 | Spleen | 22nd Nov 2022 | + | + | + | / | / |
| 9 | Stomach swab | 22nd Nov 2022 | + | + | + | / | / |
| 9 | Tonsil | 22nd Nov 2022 | + | + | + | / | / |
| 10 | Heart | 25th Nov 2022 | NEG | NEG | NEG | / | / |
| 10 | Intestine | 25th Nov 2022 | + | + | + | A/Bush<br>dog/England/058954/2022 | EPI_ISL_17521048 |
| 10 | Liver | 25th Nov 2022 | NEG | NEG | NEG | / | / |
| 10 | Lung | 25th Nov 2022 | NEG | NEG | NEG | / | / |
| 10 | Stomach | 25th Nov 2022 | NEG | NEG | NEG | / | / |
| 10 | Traches | 25th Nov 2022 | NEG | NEG | NEG | / | / |

Legend. +: positive RRT-PCR result; NEG: negative RRT-PCR result; /: no data available.

### Environmental and epidemiological investigation

To determine the extent of index site viral contamination, environmental samples were collected from eleven locations within or surrounding the bush dog enclosure (Table 3a). These had been collected shortly after the mortality event as a possible toxic / contaminant aetiology was speculated. Four matrices were investigated including water (n=12), silt (n=2), foam (n=1) and faeces (n=3). The silt, faeces and foam samples were prepared as a 10% suspension (w/v) in 0.1M PBS pH 7.2 (made locally), shaken for two minutes and incubated at room temperature for 1 hour as described previously (38). Water samples were tested without further dilution (38).

**Table 3:**
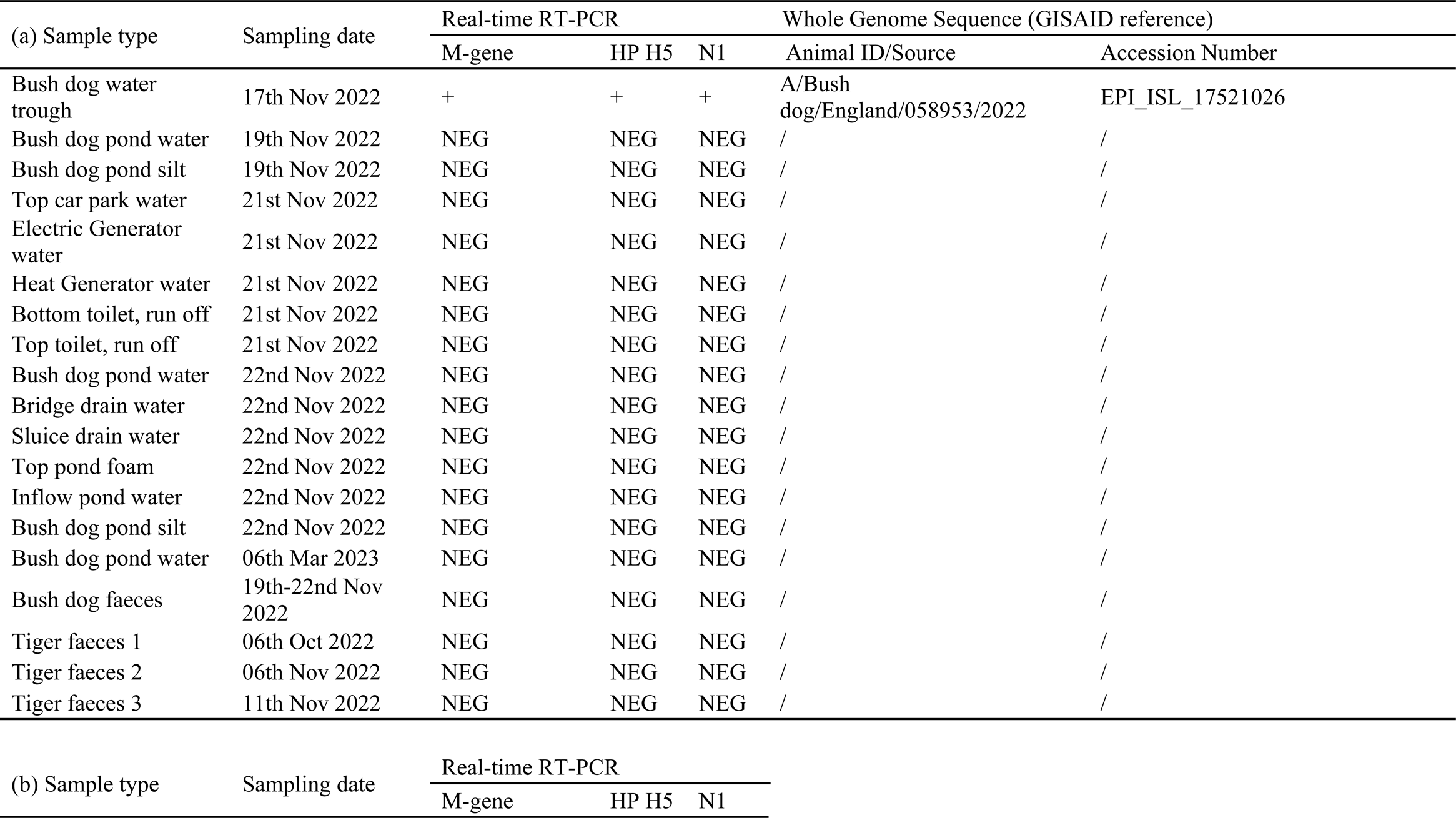
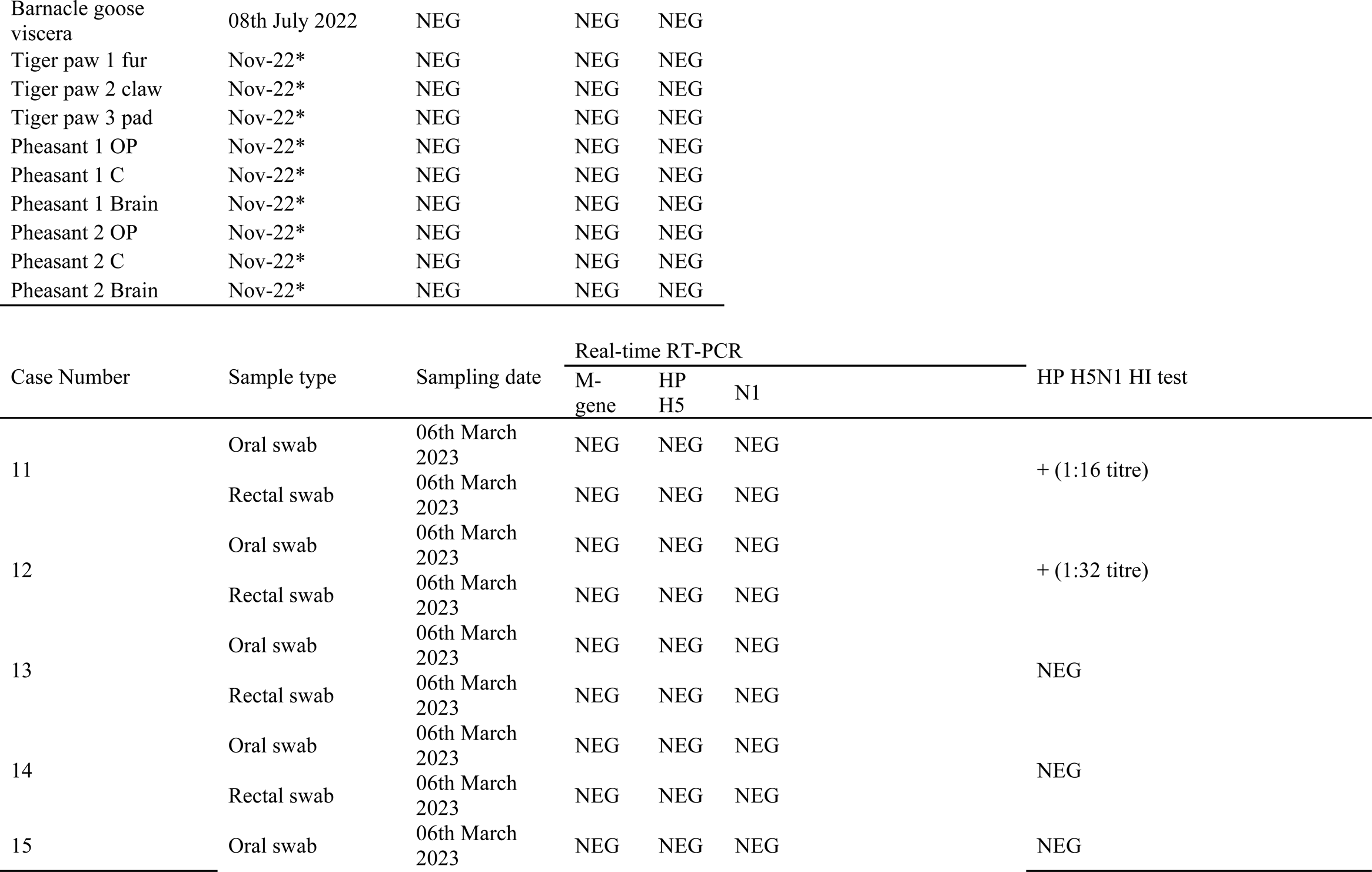
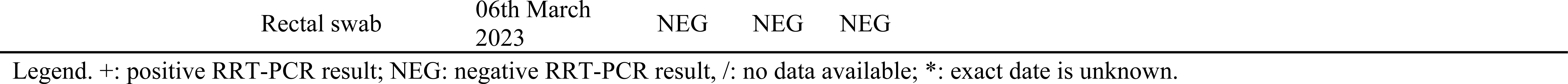
(a) Molecular assessment of enclosure environmental samples taken during the mortality event in bush dogs. (b) Molecular assessment of non-bush dog species taken during the mortality event in bush dogs. (c) Molecular and serological assessment of samples taken from surviving bush dogs.

Due to the retrospective nature of this case, samples from animals which died following outbreak in bush dogs, and still in store at the safari park, were also collected and tested (Table 3b). These samples included: An amur tiger paw from an animal which died housed near the bush dog enclosure (fur, claw and the paw pad were swabbed for this purpose) and three tiger faecal samples; swabs of oropharyngeal and cloacal cavities from two pheasants and brain tissues harvested post-mortem from the same birds. Swabs were cut into 1mL serum-free Leibovitz’s L-15 medium containing antibiotics (penicillin, streptomycin), incubated at room temperature for 10 minutes before standard viral RNA extraction. Brain tissues were prepared as a 10% (w/v) suspension in L-15 medium and incubated at room temperature for 60 minutes before using standard RNA extraction protocols (37).

Following a positive diagnosis of HPAIV infection in dogs that developed clinical disease, samples from the 5 surviving dogs were collected in March 2023 (Table 3c), including oral and rectal swabs and blood samples for molecular and serological analysis. Sera were aspirated from clotted blood samples and heat-treated at 56°C for 30 min in a water bath. All serum samples were titrated for H5-specific antibodies by using the Haemagglutination Inhibition (HI) test against four haemagglutination units of a homologous HP H5N1 antigen. Samples with a reciprocal HI titre of 16 or above were considered positive (39).

### Virological investigation

#### RNA extraction and Molecular Analysis

RNA was extracted from the environmental, tissue and swab samples using either TRIzol (Invitrogen) or the MagMAX CORE Nucleic Acid Purification Kit (ThermoFisher Scientific) using the robotic Mechanical Lysis Module (KingFisher Flex system; Life Technologies), as per manufacturer’s instructions as previously described (37). Extracted RNA was also assessed for viral nucleic acid (vRNA) using the H5 HPAIV (40) and/or NA specific (41) detection real-time RT-PCR (RRT-PCR). RRT-PCR Cq values ≤ 36.00 were considered as AIV positive. Samples with Cq >36 were considered negative. A ten-fold dilution series of titrated H5N1-21/22 HPAIV RNA was used to construct a standard curve using Agilent AriaMx software (Agilent, UK) to determine PCR efficiency which assured optimal assay performance for quantitative interpretation as previously described (37).

#### Virus isolation and propagation

For each sample, 100 µl of material was added to 100 µl of phosphate buffered saline (PBS) containing a mixture of antibiotics. The sample was incubated for 1 hour, and 100 µl was inoculated into the allantoic cavity of two specific pathogen-free (SPF) 9-day-old embryonated fowls’ eggs (EFE), as described previously (37). At 2 days post inoculation (dpi), the allantoic fluid of one EFE was collected and tested for the presence of a haemagglutinating agent using the haemagglutinin assay (HA) as previously described (37). If no HA activity was observed at 2 dpi, allantoic fluid from the remaining EFE was collected at 6 dpi and again tested by HA. HA activity >1/4 at either 2 or 6 dpi was considered positive for virus isolation. Conversely, HA activity <1/4 at both 2 and 6 dpi was considered negative for virus isolation.

#### Genomic analysis

For whole-genome sequence analysis, the extracted RNA was converted to double-stranded cDNA and sequenced using either an Illumina MiSeq or NextSeq 550 as described previously (7). Initial metagenomic analysis was performed using SeqMan NGen Software 17.5 (DNASTAR, USA) and Reference-guided application using GenBank virus reference sequences as template. Assembly of the influenza A viral genomes was performed using a custom in-house pipeline as described previously (7). All influenza sequences generated (Table S1) and used in this study are available through the GISAID EpiFlu Database (https://www.gisaid.org). Initial analyses sought to compare the sequences obtained from the Bush dogs and associated samples with contemporary H5N1 sequences from the UK and Europe. To achieve this, all avian H5N1 clade 2.3.4.4b sequences containing all eight influenza gene segments available on the GISAID EpiFlu platform between 1 January 2020 and 17 July 2023 were downloaded. Duplicate sequences were removed before all sequences were aligned on a per segment basis using MAFFT v7.520 (42). The alignments were visually inspected using AliView version 1.26 (43) and sequences which did not cover the entire open reading frame (ORF) across all eight gene segments were removed. The remaining sequences were aligned and trimmed to the ORF for each segment before generating a concatenated alignment using SeqKit (44) and then used to a infer maximum-likelihood phylogenetic tree using IQ-Tree version 2.2.3 (45). This resultant phylogenetic tree contained over 2,000 sequences and was therefore sub-sampled to cover 98% of the diversity within using PARNAS (46), which reduce the dataset down approximately 300 sequences whilst still containing representatives of the predominant UK genotypes. The sub-sampled dataset was then used to infer maximum-likelihood phylogenies for each gene segment using IQ-Tree along with ModelFinder (47) and 1,000 ultrafast bootstraps (48). Once the genotype of the Bush dogs and associated sequences was determined, all sequences of the relevant genotype were retrieved from the original dataset (prior to sub-sampling) and used to generate time-resolved phylogenies using TreeTime (49). Phylogenetic trees were visualised as described previously (7). Nucleotide identity between sequences was determined as described previously (50).

## Results

### Clinical Setting

The bush dogs had been clinically healthy with no significant clinical concerns in the preceding 3 years. A timeline of disease status is presented in Figure 1. Case 1, day 1, male, 9-year-old bush dog was found dead adjacent to the entrance of the nest box with a CCTV review showing hindlimb ataxia, forelimb hypermetria, depression and polyuria. Case 2, day 1, male, 3-year-old was moribund and assessed under anaesthesia with uraemia, raised creatinine, hyperphosphataemia, total bilirubinaemia, and a raised alanine aminotransferase was noted, and was found dead the following morning, day 2. Case 3, day 2, 4-year-old female was found dead in the main pond within the exhibit. Case 4, day 3, a 5-year-old female was euthanised due to severe ataxia combined with depressed responsiveness and polyuria. Case 5, day 3, a 4-year-old male was euthanized due to severe ataxia combined with depressed responsiveness and polyuria. Case 6, day 4, a 9-year-old female presented with severe hind-limb ataxia, weakness, reduced activity and polyuria, and was euthanized. Case 7, day 5, a 4-year-old female appeared clinically healthy during the morning assessment but within 2 hours was moribund and unresponsive and was euthanized on welfare grounds. Case 8, day 5, a 3-year-old male was clinically normal at the morning assessment but within 2 hours had developed severe ataxia and generalised weakness, vocalising and was euthanized on welfare grounds. Case 9, day 6, a 1-year-old female was found dead during the morning check. Case 10, day 9, a 4-year-old male presented with severe hind limb weakness, knuckling, and ataxia, and was euthanized on welfare grounds. The remaining five animals, two males and three females, age range 18 months to 4 years were clinically normal during the same period and remained unaffected from the event.

**Figure 1:**
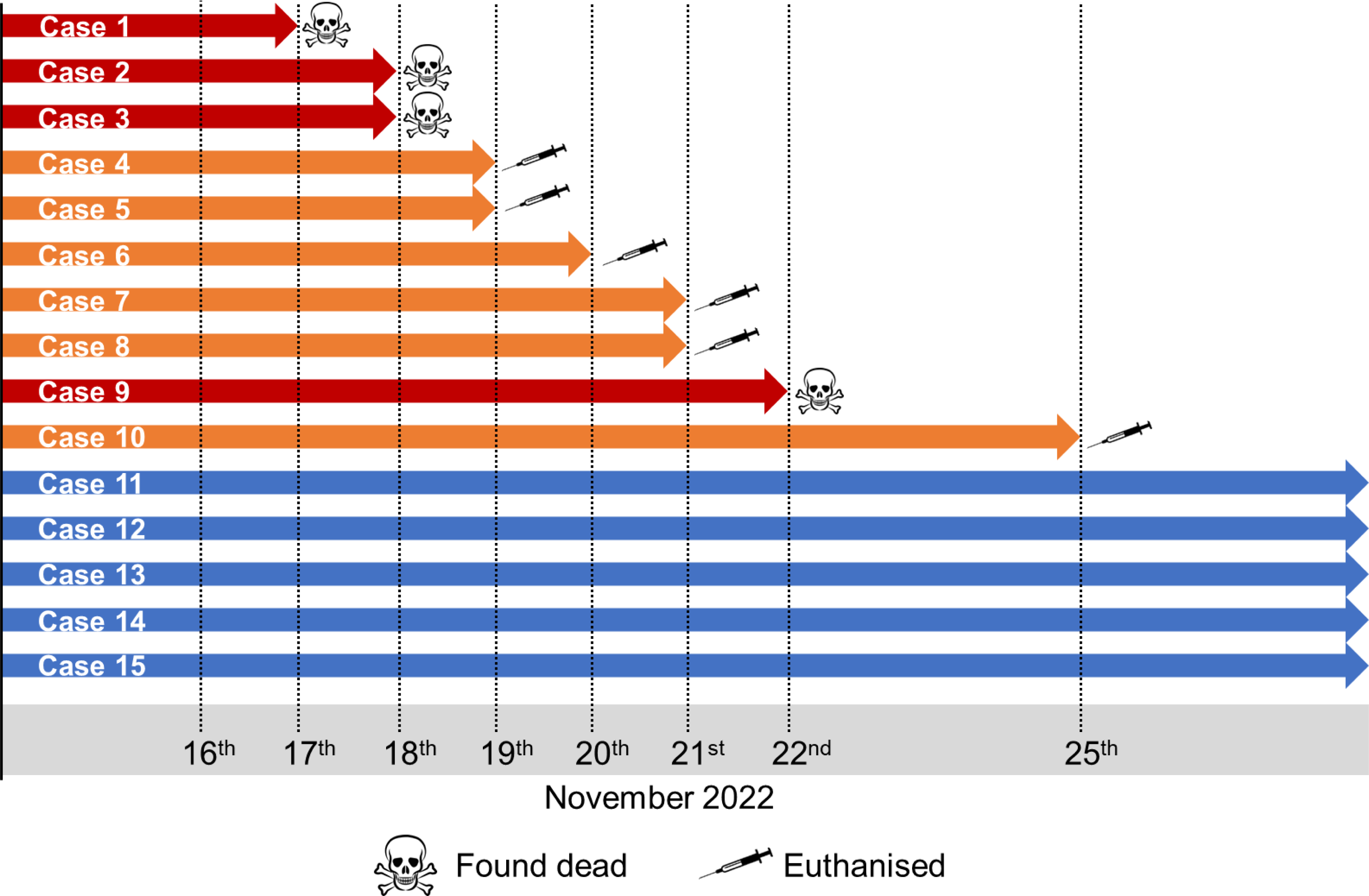
Timeline of mortality event within the bush dog enclosure

Pathological investigation and preliminary collateral analysis were performed at private laboratories. On gross examination, the primary findings were diffuse mild to moderate hepatomegaly (8/10), with multifocal to diffuse pale tan discolouration within liver lobes (7/10) and mild to moderate ascites (7/10). Intracardiac euthanasia made gross assessment of the lungs challenging in six cases, but mild to moderate multifocal pulmonary haemorrhage was seen in three of the remaining four cases. Other less common gross findings included bilateral adrenomegaly (6/10), diffuse splenomegaly (3/10), and segmental small intestinal congestion and haemorrhage (4/10).

Microscopically, within many of the tissues examined, vascular changes were observed, ranging from occasional subacute multifocal moderate to marked fibrinoid necrosis of small arterioles without associated inflammation, to acute, multifocal, mild to severe necrotising and sometimes leukocytoclastic phlebitis of vessel walls, with fibrin thrombi, and necrosis of adjacent structures. In the liver, random mild to moderate acute multifocal necrosis (Figure 2a) and inflammation affecting both the parenchyma, and blood vessels and bile ducts within portal triads, was seen in all ten cases. Foci of hepatocellular necrosis were rarely surrounded by minimal neutrophilic and histiocytic inflammation.

**Figure 2:**
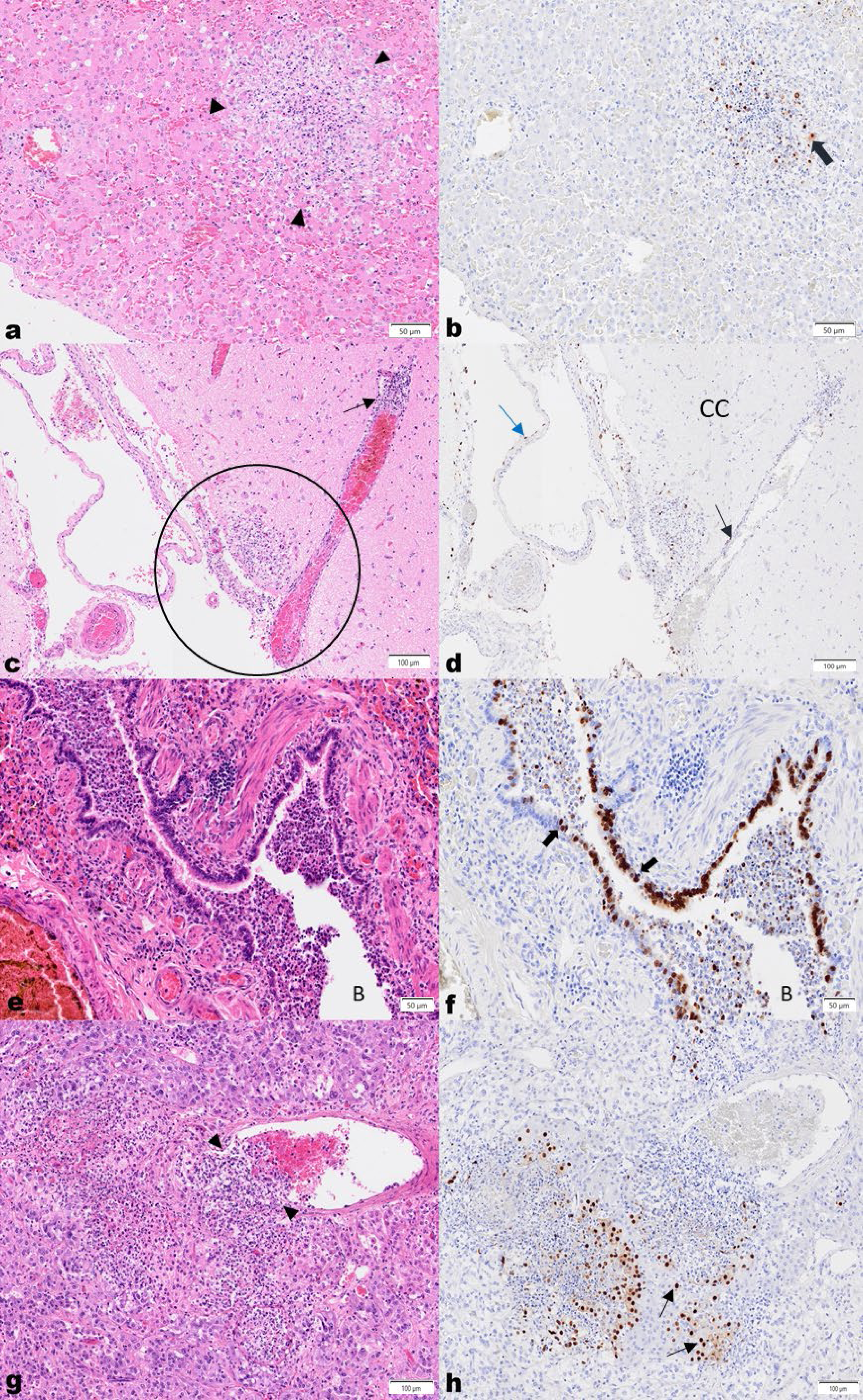
High pathogenicity avian influenza virus H5N1 in bush dogs. (**a**) Liver, H&E. Acute, random focal hepatocellular necrosis, comprised of karyorrhectic debris and fibrin exudation with rare neutrophils and histiocytes. (**b**) IHC shows viral antigen in hepatocytes (thick brown arrow) on the periphery of the lesion, and in cellular debris. (**c**) Brain, cerebral cortex (**CC**), H&E. Vasculitis with segmental multifocal infiltration of a venule wall with neutrophils and histiocytes, is shown (thin black arrows). Multifocal to locally extensive necrosis and inflammation extends from the meninges into the adjacent neuropil (black circle). (**d)** IHC demonstrates antigen in vascular endothelial (black arrow) and arachnoid epithelial cells (blue arrow). (**e**) Lung, H&E. The bronchiolar lumen (**B**) is filled with degenerate and viable neutrophils, macrophages, eosinophilic material and shed epithelial cells, and the lamina propria and submucosa of the bronchiolar wall is diffusely mildly infiltrated by lymphocytes, histiocytes and fewer neutrophils. Adjacent alveolar spaces are collapsed and alveolar septae congested and mildly expanded by neutrophils and macrophages. (**f**) IHC labelling is present in bronchiolar epithelial cells (black arrows) and within shed epithelial cells and macrophages in the lumen. (**g**) Adrenal gland, H&E. A subacute focal severe necrotising vasculitis is seen with expansion of the vascular wall with neutrophils, pyknotic cellular debris and fewer mononuclear cells (black arrowheads), with subacute multifocal severe necrosis and inflammation of the adjacent epithelial cells in the acini of the adrenal medulla and cortex. (**h**) IHC highlights viral antigen in the acinar epithelial cells (black arrows).

Variable changes in the lungs were present in six bush dogs. The bronchioles and adjacent alveoli were predominantly involved in four cases where the bronchiolar lumina were filled with degenerate neutrophils and epithelial cells with attenuation and necrosis of bronchiolar epithelium. (Figure 2; e). Mild neutrophilic, lymphocytic and histiocytic infiltration of the lamina propria and submucosa was also seen. Adjacent to the affected airways, alveoli were variably filled with fibrin, oedema, alveolar macrophages, neutrophils and erythrocytes. Less commonly, a more interstitial pattern was seen with mild to moderate expansion of the alveolar septae with macrophages, lymphocytes and neutrophils. Occasionally, vascular pathology was present, characterised by segmental infiltration of the vascular wall with leucocytes and angiocentric necrosis.

The brain was collected in four cases, and in all, the leptomeninges were expanded by mild to marked multifocal subacute inflammation. In three cases (7, 9 and 10), the infiltrate was composed mainly of viable and degenerate neutrophils, with rarefication and inflammation of the adjacent neuropil (Figure 2: c). In the fourth case (6), the infiltrate consisted predominantly of lymphocytes, macrophages, and plasma cells with fewer neutrophils, and locally extensive involvement of the choroid plexus on the section examined. In all cases, randomly within the cerebral cortex and brainstem, and rarely in the cerebellum, few to multiple foci of mild to moderate necrosis and inflammation consisting mainly of aggregates of macrophages and glial cells, necrotic debris, and rare neuronal necrosis, were seen. Grey matter and white matter were both affected. In the neuropil, vascular changes were mild, and characterised by an acute multifocal vasculitis with infrequent mild histiocytic perivascular cuffing. However, in the leptomeninges, vascular lesions ranged from a mild to severe, and acute to subacute multifocal necrotising vasculitis (Figure 2: c).

In three cases, multifocal marked acute cortical and medullary epithelial necrosis with neutrophilic infiltration, congestion, and focal subacute severe leukocytoclastic vasculitis (Figure 2: g and h) was observed in the adrenal glands.

In lymph nodes examined from three bush dogs, findings varied from moderate multifocal lymphadenitis with predominantly subcapsular sinus neutrophilia and histiocytosis, and multifocal necrosis, to severe, diffuse acute lymphadenitis. Severe changes were characterised by expansion of the sinuses with viable and degenerate neutrophils, histiocytes and fewer erythrocytes, and multifocal necrosis of the cortex and paracortex with infiltration of large numbers of degenerate neutrophils.

Within adjacent small blood vessels, the walls were multifocally infiltrated by degenerate neutrophils and karyorrhectic debris, and in the lumina, multifocal thrombi comprised of fibrin, macrophages and neutrophils were present.

The spleen was minimally affected, but in all four reported cases of mild acute splenitis, multifocal clusters of viable and degenerate neutrophils and occasionally karyorrhectic debris and histiocytes, were seen in the white pulp. Haematopoietic precursors (extramedullary haematopoiesis) and haemosiderin laden macrophages were scattered within the red pulp. No vascular changes were apparent.

In the intestinal sections examined, inflammation was mostly mild, multifocal, diffuse and chronic. However, in the small intestine, evidence of mild to moderate active inflammation and necrosis was noted sporadically, with crypt abscessation, and foci of necrosis and monocytic inflammation within and between the inner circular and outer longitudinal smooth muscle layers, often associated with capillaries and in the submucosal and myenteric plexuses. In one case, acute, moderate multifocal necrotising typhlitis and colitis was identified, with lymphoid depletion and mild acute multifocal vasculitis rarely present in the small vessels of the submucosa.

Where the eye was sampled, significant widespread and occasionally severe inflammation was found centred around vessels, with resultant anterior uveitis, optic neuritis and infrequently, retinitis. Multifocal necrotising neutrophilic inflammation of the third eyelid gland (nictitans gland) was captured in one section.

Interestingly, the heart (three cases) and pancreas (two cases) appeared mildly affected in the sections examined, displaying mild multifocal acute degeneration and necrosis with minimal associated inflammation. Incidental mild, chronic, multifocal interstitial inflammation was reported in the kidneys but a mild to moderate multifocal acute vasculitis was present in some sections. On the serosal surface of many abdominal organs, particularly the liver, a mild to moderate acute multifocal to diffuse fibrinous peritonitis was seen. A detailed summary of the gross and histopathology reports provided by the park owner and the local specialist laboratory is provided in the supplemental material (Supplementary Table 1).

The changes described were suggestive of a severe acute systemic infection, which was initially thought to be bacterial, but a viral or parasitic aetiology could not be ruled out. Collateral analysis was carried out to exclude common causes of death. Secondary or contaminant bacteria were isolated from affected organs, such as *Escherichia coli, Staphyloccus aureus, Clostridium perfringes* and *Clostridium sordelli*, *Haemophilus parainfluenzae* and *Proteus* spp. Molecular tests (PCRs) performed at a private laboratory excluded presence of SARS-CoV-2, Leptospirosis, RHDV type 1 and Canine Adenovirus. Toxicology analysis for Ethylene glycol were negative. Despite this further ancillary testing, no aetiologic agent was identified and preliminary clinical investigation was inconclusive.

### Immunohistochemistry

Immunohistochemistry was performed using an anti-influenza A antibody targeting viral nucleoprotein, on formalin fixed paraffin embedded tissues previously submitted for routine histopathologic analysis (Table 1). Generally, infrequent viral antigen was detected in macrophages and in cellular debris within and on the periphery of necrosis and inflammation in the many affected organs. Angiocentric necrosis and inflammation was sometimes accompanied by sporadic labelling of the vessel walls, particularly the endothelium and tunica media.

More specifically, within organs, occasional viral antigen was present in hepatocytes (Figure 2b), and in bile duct epithelium in the liver. In the brain, viral antigen was abundant in ependymal cells lining the lateral and third ventricles, and less frequently, multifocally in neurons, axons, and glial cells in the neuropil. Labelling was also present in the meningeal epithelial cells in the subarachnoid space (Figure 2: d). Moderate sporadic labelling was seen in bronchiolar epithelium (Figure 2f) but was minimally visible in the alveolar septae.

Significant immunolabelling was observed in acinar cells of the adrenal cortex (zona fasciculata and reticularis), and less commonly in cells of the adrenal medulla (Figure 2h). In the eye, viral antigen was detected in macrophages around vessels and within necrotic debris, particularly in the iris, uvea and meninges of the optic nerve. Moderate viral antigen was found in the acinar cells of the nictitans gland of the eye. Strong labelling was seen in the nuclei of smooth muscle cells of one or both the smooth muscle layers of the small and large intestines, but infrequently within macrophages, in the vascular endothelium of the capillaries, and in ganglion cells of the plexuses between them. Rarely, viral antigen was visible in mucosal epithelial cells of the stomach, and enterocytes in the small and large intestines. Occasional viral antigen was present in the serosal epithelium on the surfaces of the intestines, abdominal organs, and urinary bladder. Minimal antigen was rarely seen in cardiomyocytes.

#### Virological assessment

Viral RNA (vRNA) was detected across a broad range of tissues sampled (Table 1) in all bush dog cases. Due to the retrospective nature of the detection, and the working practices of the team on location during the disease event, a significant number of tissues were collected at PME and these were assessed following retrospective confirmation of influenza A virus of avian origin as the causative agent. All samples tested positive across the suite of molecular tests used to detect the influenza A: M-gene, the high pathogenicity H5 haemagglutinin (H5 HP) and the N1 neuraminidase gene. Exceptions to this were oral and rectal swabs from case 6 and the intestine from case 10, all of which tested negative across all three molecular assays. Of significant interest were the positive results obtained from a urine sample that was aspirated directly from the bladder during the PME of case 5. Virus isolation and propagation was successful, resulting in the harvesting of one viral isolate per dog.

### Environmental and epidemiological investigation samples

During the mortality events, staff had also collected a suite of environmental samples (Table 3) from their enclosure which could be assessed for evidence of environmental contamination. An additional sample set included samples from animals at the park that were considered at risk from the agent, and which died or were euthanised during or shortly after the canid mortality event. Interestingly, from all these samples, only one sample of water taken from a drinking trough within the bush dog enclosure tested positive for viral RNA although an isolate was not successfully recovered from the sample (Table 3). Samples from the five surviving dogs, collected approximately four months after the disease event, were also negative at RRT-PCR. However, mild seroconversion was detected in 2 out of 5 dogs following HI test, demonstrating exposure to virus or viral antigen.

### Genomic analyses

Initially, samples across Cases 6, 8 and 9 were subjected to metagenomic analyses; this included liver, kidney and brain samples from case 6 and three tissue pools of liver, kidney and spleen from cases 8 and 9. The analysis revealed sequence reads matching those of influenza A virus. This initial metagenomic findings in combination with the RRT-PCR results prompted further investigation of the samples obtained from the Bush dogs and WGS was attempted on tissue samples or viral isolates from the deceased animals and the positive environmental sample, resulting in 11 full H5N1 HPAIV genomes being sequenced. The 11 genomes demonstrated 99.4-100% nucleotide identity to each other across all eight influenza virus gene segments. Initial phylogenetic analysis revealed that the genomes obtained from the bush dog samples clustered with those of the UK AIV09 genotype (7) (also known as the AB genotype according to the EU Reference Laboratory schema) (Figure S1), which was the predominant genotype in the UK from October until May 2023. To further investigate potential incursion routes for H5N1 into the captive Bush dog population, 10 of the 11 genomes from the safari park were concentrated and combined with a more fulsome dataset containing only genomes of the AIV09 (AB) genotype which were then used to infer time-resolved phylogenies. The omitted sequence (A/Bush dog/England/037675/2022) contained a number of gaps and was removed from these subsequent analyses. From these analyses, it was demonstrated that the bush dog genomes showed high similarity with AIV09 (AB) H5N1 HPAIV sequences from the UK, but formed a distinct group (Figure 3a), with a time to most common recent ancestor (tMRCA) estimate of 24^th^ September 2022 (range: 9^th^ September 2022 to 13^th^ October 2022). Within this group, the 10 bush dog sequences were divided into two separate sub-groups. Sub-group A contained the environmental sequence generated from the water trough in the Bush dog enclosure (A/Environment/England/058953/2022), along with two sequences from Bush dogs (A/Bush dog/England/037682/2022 (case 3) and A/Bush dog/058954/2022 (case 10)) and were generated from samples collected from the 17^th^ ^of^ November through to the 25^th^ of November, covering the entire sampling period. Sub-group B contained the remaining seven Bush dog sequences, generated from samples collected between the 18^th^ ^of^ November and 22^nd^ November. When looking at the amino acid changes across the bush dog sequences, 10 variable sites were observed across six viral proteins: polymerase basic protein 1 (PB1) and 2 (PB2), polymerase acidic protein (PA), HA, matrix protein 2 (M2) and non-structural protein 1 (NS1) (Figure 3B). The PB2 protein contained the most variant sites (five), which included changes at positions 627 and 701 which have both been reported to be involved in adaptation of AIVs to mammalian hosts (27). Interestingly, whilst nine of the 10 bush dog sequences, including the environmental sample, possessed a lysine (K) at position 627, one sequence (A/Bush dog/England/068984/2022 (case 4)) did not. However, this sequence did contain an asparagine (N) at position 701, whereas the other sequences possessed an aspartic acid (D). When comparing the phylogenetic and amino acid variability, it was found that sub-groups A and B were distinguished by three variable sites in the PB2 (position 403 and 598) and PA (position 531) proteins. Within sub-group B, there were also three sequences (A/Bush dog/England/058198/2022, A/Bush dog/England/037689/2022 and A/Bush dog/England/037690/2022) that showed different amino acid substitution from the other sequences in sub-group B (PB2 position 123 and NS1 position 183), however, there were also other changes seen within this sub-group at different positions.

**Figure 3.**
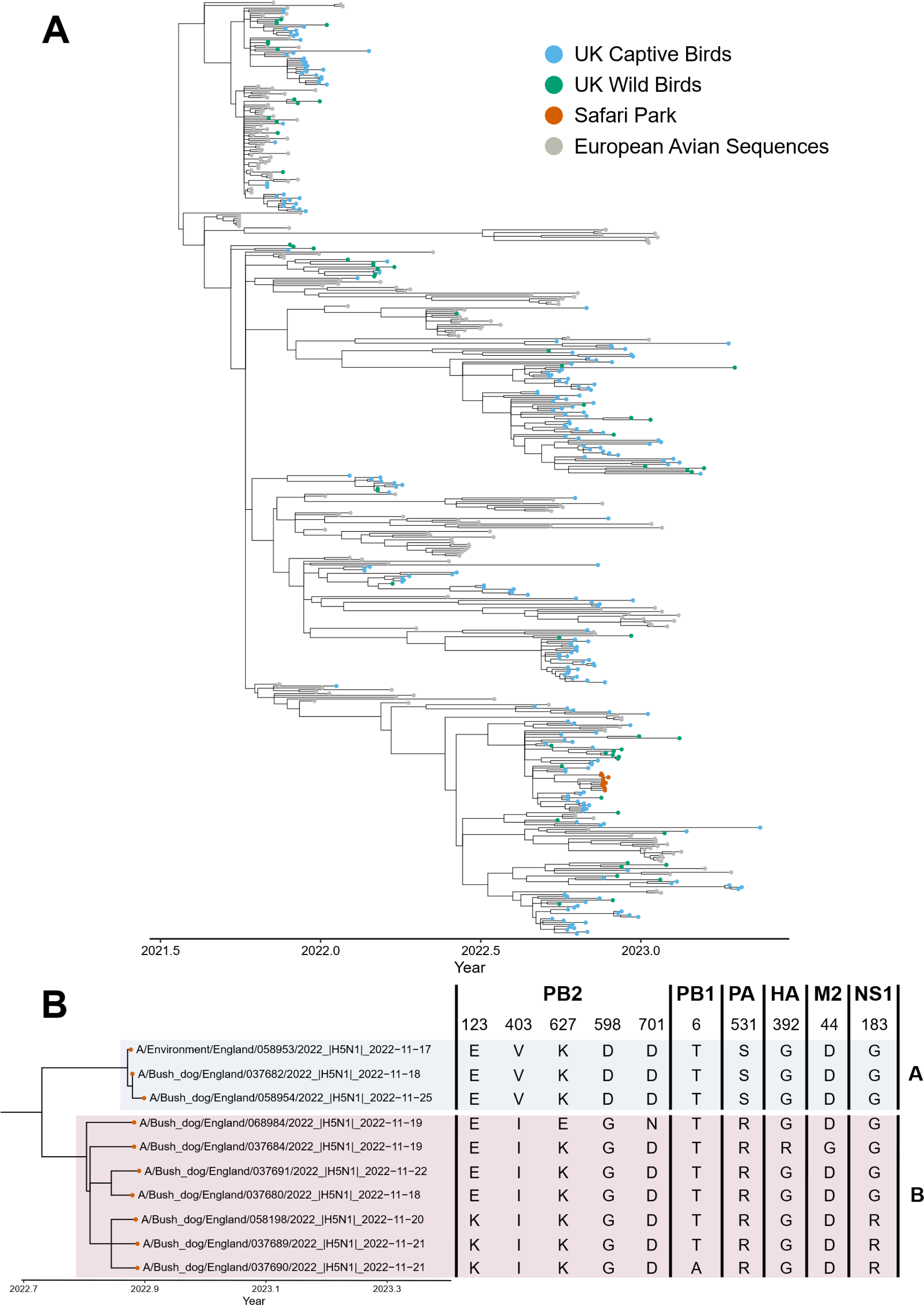
**(a)** Time-resolved phylogenetic analysis of H5N1 HPAIV genotype AIV09 (AB) concatenated from genomes. Sequences from UK captive and wild birds, as well as the sequences obtained from the bush dogs are coloured accordingly. **(b)** A subset of the phylogenetic tree shown in A focussing on the bush dog sequences and the amino acid changes therein. Sub-groups A and B described in the text are highlighted.

## Discussion

The detection of influenza A virus of avian origin H5N1 as the causative agent of mortality in bush dogs was an unusual and unexpected event. Influenza A virus infection was not on the list of differentials for causative agent in this disease event and was initially only detected from sequence analysis following next generation sequencing of submitted samples. Sequence reads for influenza A virus led to the described diagnostic pathway that conclusively defined clade 2.3.4.4b H5N1 influenza A virus of avian origin as the cause of disease. The internal distribution of the virus demonstrated systemic and wide-spread infection affecting multiple organs and clearly demonstrate that HPAI was the primary cause of death. Viral excretion through the respiratory and intestinal tract could be speculated although swabs (oral and rectal) collected from case 6 did not detect virus. The interesting detection of HP H5N1 vRNA from case 5 from a urine sample may hint at possible horizontal transmission routes although there is no further evidence to substantiate this hypothesis as an unexpected shedding pathway for an Avian Influenza virus.

Histopathologic findings demonstrated a severe acute systemic disease characterised by vasculitis, and widespread necrosis and inflammation in many organs, specifically the liver, brain, lung, and adrenal glands. Immunohistochemistry confirmed the presence of viral antigen most often seen in the brain, lung, adrenal glands, lymph nodes and liver, but also within vascular walls. The brain and lung are common targets in spill over mammalian HPAI H5N1 infections (5, 51–53), with virus potentially gaining entry following inhalation into the respiratory tract. Other routes, such as infection through the intestinal barrier into the vascular system, following ingestion of infected bird carcasses have also been established as a route of H5N1 infection in cats, with subsequent viraemia (54). In these bush dogs, both lung and intestinal lesions were seen, with antigen detected in both tracts, although generally mild, supporting a multifactorial pathogenesis. Dissemination of the virus to the brain haematogenously with penetration of the blood-brain-barrier into the cerebrospinal fluid (CSF) has been proposed (55) and would account for virus detected in the meninges and spread to the adjacent neuropil. Vasculitis, and in particular, phlebitis was a significant feature of infection in this species but is infrequently described in mammalian HPAI H5N1 infection. Recent reports in red foxes (10, 51) and in historical and recent cases of HPAI H5N1 in naturally infected cats (55, 56) report endotheliotropic behaviour of the virus, with endothelial damage and vasculitis in seen in multiple organs, which resembles the endotheliotropism seen in the pathogenesis of HPAI H5N1 infections in terrestrial poultry (54). However, leukocytoclastic vasculitis, defined as a small-vessel neutrophilic vasculitis with fragmented nuclei present (57) has only recently been described in naturally infected cats (56).

The exposure route to influenza A virus of avian origin in this case is hard to conclusively define. The bush dogs had been fed a diet that included frozen shot wild birds and game. In the absence of local disease events that may have been transferred to the bush dogs in the enclosure, infection through ingestion of infected meat / offal would appear to be the most likely route of infection. Another potential infection route is through scavenging of any wild bird carcases/any sick wild birds landing in the un-netted pen. Other routes of infection including indirect contact (e.g., wild bird faeces) are possible but less likely and would not fit with the rapid onset of infection across a number of dogs within a short time frame. Wild bird activity was observed on the site during epidemiological investigations and black headed gulls, greylag geese, pink footed geese as well as corvids, pigeons and pheasants had been observed in the vicinity. The shot wild game supplied as food had originated from the neighbouring shooting estate but no reports of clinical signs or die-offs in avian species had been reported. Further, the stream/pond in the bush dog enclosure was supplied with water from a reservoir upstream that was frequented by wild bird species, although a sluice was in place to prevent carcasses being washed down from the reservoir and the water was relatively fast flowing, again making this an unlikely infection route. Critically, the rapid development of disease and acute clinical outcome suggests that the dogs received a high dose of infectious material and so ingestion of material must be considered the most likely route of infection. Genetic assessment from tissues positive for each animal was undertaken to try and evaluate the possibility of dog-to-dog transfer. Local atmospheric conditions are known to influence environmental virus survival with lower temperatures promoting persistence (38, 58). In early October 2022 the day-night high-low temperature range near this site was 17 to 11 °C. For samples collected on 17 to 22 November 2022 and 06 March 2023, day-night readings were consistent at 8 – 11 °C and 4 – 8 °C respectively (59), hence favourable to virus survival. Other factors can impact detection of environmental viral RNA from water, dilution (rain), volume (lake) and flowrate (river) (58). This might account for a single M-gene signal detection from the bush dog water trough (17/11/2022), a fixed volume container of still water (Table 3a). All bush dogs had access to the water trough which we found contained viral RNA. Since five bush dogs survived this lowers the likelihood of water being the route or main source of virus introduction and trough contamination by infected bush dogs being a plausible explanation for this vRNA detection.

The genetics of the virus demonstrated a high level of viral genetic homogeneity across all eight viral segments of the sequences from the bush dogs. Phylogenetic analysis demonstrated that these sequences were consistent with the AIV09 (EURL: AB) H5N1 HPAIV genotype which predominated in the UK during the 2022-2023 autumn/winter period (7). Time-resolved phylogenetic and amino acid analysis found that all the sequences from the bush dogs were the result of a single introduction, however whilst there were amino acid substitutions, these do not appear to have been consistently maintained. Taken together, this suggests that transfer between dogs is unlikely, and that a common source of infection is responsible, although it is impossible to definitively conclude whether dog-to-dog transmission occurred. Critically, of the original bush dog population within the enclosure, 5 animals survived remaining clinically normal throughout. This may indicate that these dogs had not received a dose of virus sufficient to drive a productive infection. This is further supported by low level serological responses being detected in two of the animals (cases 11 and 12) that may indicate exposure to antigen or a low-level infection that was cleared by the host immune response. From the perspective of zoonotic risk, the well-established marker of mammalian adaptation (E627K) was detected in all but one of the bush dog sequences generated. This mutation alone is insufficient to drive an increase in zoonotic risk and so the risk to human populations must be considered very low.

Infection of unusual species with influenza A of avian origin H5N1 raises important questions about the potential implications for infection of conservation species. Clearly, the increasing detection of mammalian infection with these viruses is of global interest, not least through the potential for mammalian adaptation and establishment of mammal-to-mammal transmission. This factor is being monitored globally wherever mammalian infection is detected, and potential adaptive mutations scored for relevance to potential viral adaptation. Critically, understanding the implications of infection of captive, potentially rare or endangered species is key to enabling prevention of such occurrences. Clearly the feeding of wild shot birds to captive carnivores whilst infection pressure is high in wild birds should be discouraged in line with similar recommendations given to keepers of birds of prey or in alternative a rigorous risk assessment should be carried out before any carcass is fed to any of these animals.

## Acknowledgements

MF, SMR, AMPB, SMR, NM, CJW, SST, JJ and ACB were part funded by the UK Department for the Environment, Food and Rural Affairs (Defra) and the devolved Scottish and Welsh governments under grants SE2213, SE2227, SV3400 and SV3006. ACB and JJ were also part funded by the Biotechnology and Biological Sciences Research Council (BBSRC) and Department for Environment, Food and Rural Affairs (Defra, UK) research initiatives ‘FluMAP’ [grant number BB/X006204/1] and ‘FluTrailMap’ [grant number BB/Y007271/1], and the Medical Research Council (MRC) and Defra research initiative ‘FluTrailMap-One Health’ [grant number MR/Y03368X/1] as well as by Federal funds from the National Institute of Allergy and Infectious Diseases, National Institutes of Health, Department of Health and Human Services (USA), under contract no. 75N93021C00015. This investigation was also partly funded by APHA GAP-DC SE0565 project. We thank the scientific and support staff of the Pathology and Virology departments of APHA Weybridge and of the Surveillance and Laboratory Services department of the APHA Veterinary Investigation Centres. We acknowledge the authors, originating and submitting laboratories of the sequences from GISAID’s EpiFlu Database on which this research is based, Veterinary Pathology Group and PALS for their histopathology expertise in initial reporting, and analyses described in text. All submitters of the data may be contacted directly via the GISAID website (www.gisaid.org).

---

**Figure S1.**
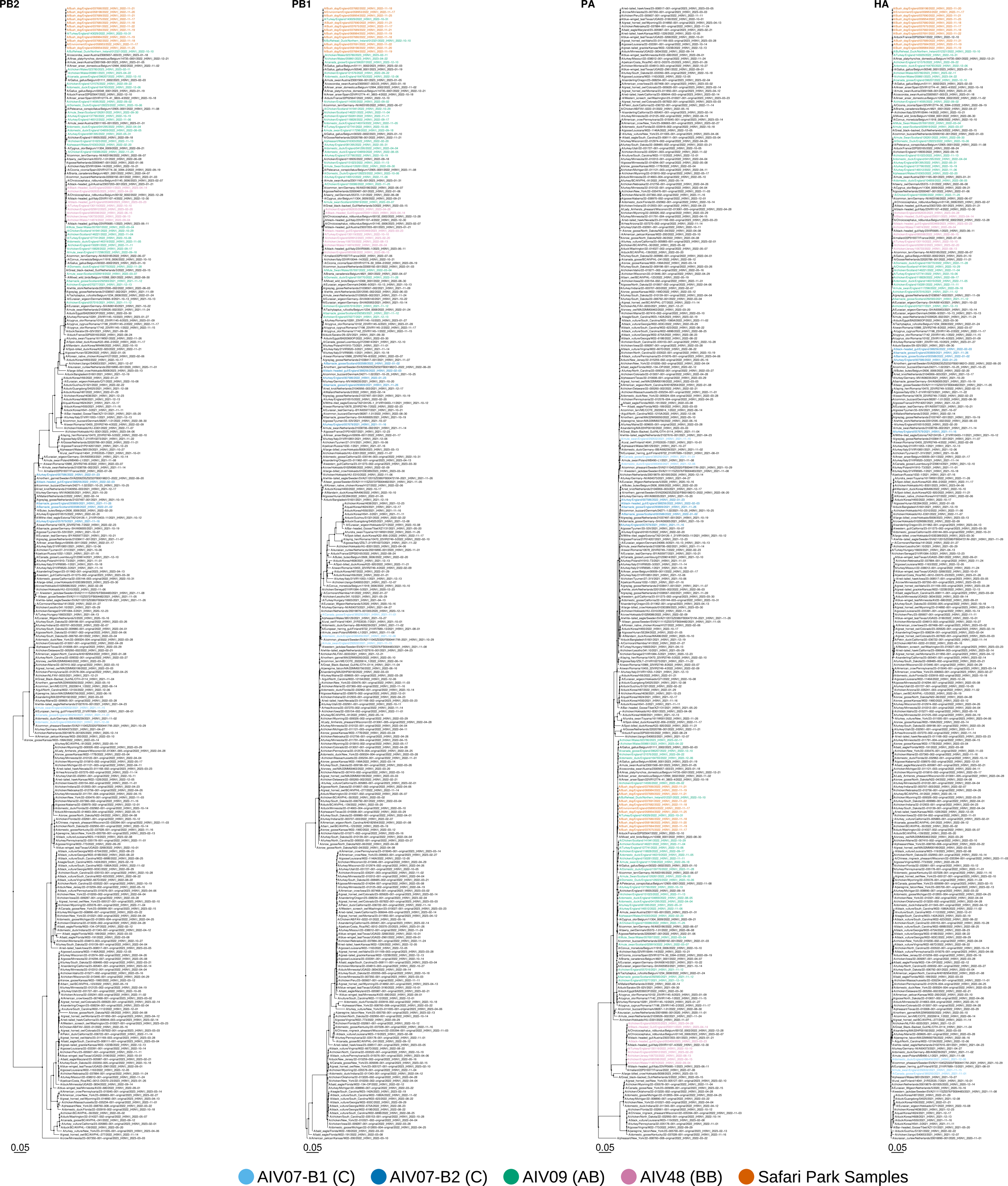
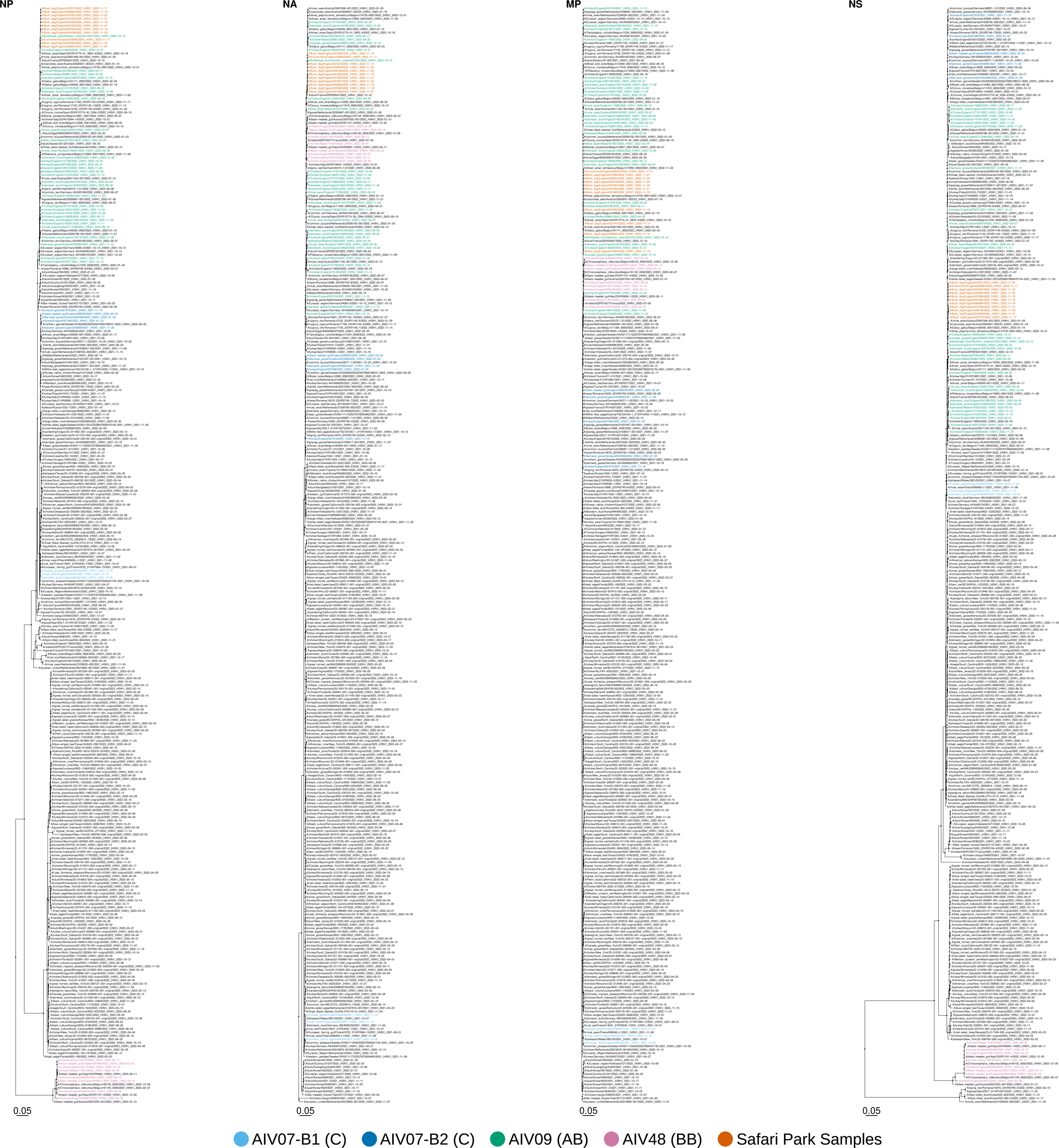
Midpoint-rooted maximum-likelihood phylogenetic trees containing the sub-sampled set of H5N1 clade 2.3.4.4b global sequences along with those obtained from the Safari park. Sequences representative of the predominant genotypes detected within the UK (along with their European Union Reference Laboratory genotype designation) are coloured accordingly.

